# Integrating population-level and cell-based signatures for drug repositioning

**DOI:** 10.1101/2023.10.25.564079

**Authors:** Chunfeng He, Yue Xu, Yuan Zhou, Jiayao Fan, Chunxiao Cheng, Ran Meng, Lang Wu, Ruiyuan Pan, Ravi V. Shah, Eric R. Gamazon, Dan Zhou

## Abstract

Drug repositioning presents a streamlined and cost-efficient way to expand the range of therapeutic possibilities. Drugs with human genetic evidence are more likely to advance successfully through clinical trials towards FDA approval. Single gene-based drug repositioning methods have been implemented, but approaches leveraging a broad spectrum of molecular signatures remain underexplored. We propose a framework called “TReD” (Transcriptome-informed Reversal Distance) that integrates population-level disease signatures robust to reverse causality and cell-based, drug-induced transcriptome response profiles. TReD embeds the disease signature and drug response profiles in a high-dimensional normed space, quantifying the reversal potential of candidate drugs in a disease-related cell-based screening. Here, we implemented this framework to identify potential therapeutics relevant to COVID-19 and type 2 diabetes (T2D). For COVID-19, we identified 36 drugs showing potential reversal roles. Notably, nearly 70% (25/36) of the drugs have been linked to COVID-19 from other studies, with seven drugs supported by ongoing/completed clinical trials. For T2D, we observed reversal signals for 16 compounds on multiple disease signatures. Five drugs are supported by published literature, affirming potential therapeutic value. In summary, we propose a comprehensive genetics-anchored framework integrating population-level signatures and cell-based screening that has the potential to accelerate the search for new therapeutic strategies.

## Introduction

Conventional drug development is expensive in cost and time, spanning over a decade with substantial financial investment to go from target identification to therapeutics(DiMasi et al., 2003); Wouters et al. (2020). Drug repositioning represents a cost-effective strategy to extend the utility of existing drugs, leveraging previous studies on safety in preclinical and clinical studies to accelerate drug application.

Genetic disease associations may play a significant role in repositioning, since empirical data suggests that drug targets with genetic support are 2.6 times greater to be successful(Minikel et al., 2024). Genome-wide, transcriptome-wide, and proteome-wide association studies (GWAS, TWAS, PWAS) are powerful genetic tools to predict drug repurposing candidates: for example, the transcriptome-wide significant association between *TYK2* and critical illness in COVID-19(Pairo-Castineira et al., 2021), facilitated the drug repurposing of baricitinib, which has demonstrated therapeutic benefit in subsequent clinical trials(Rubin, 2022).

However, drug repositioning focused on a single gene target may not be sufficient to elicit a meaningful therapeutic response as complex diseases involve intricate networks of pathways and interactions(Pushpakom et al., 2019). The growing evidence for polypharmacology calls for the adoption of experimental-computational multi-target approaches(Paolini et al., 2006). Most existing approaches rely on differential gene expression (DGE) to generate a list of prioritized genes. However, the transcriptome perturbations identified by DGE are challenging to interpret as molecular consequences or causes of disease, with one recent study claiming a higher likelihood of implicated genes being of the former class(Porcu et al., 2021). Moreover, methodologies grounded in DGE analysis inherently struggle to determine the direction of pharmacological effect (therapeutic or harmful) on disease when lacking the context of a known disease pathogenesis.

In contrast, a method that performs an inverse pattern-matching alignment of disease signatures and drug response profiles(Chen et al., 2017) suggests that, for a given disease signature comprising genes with altered up/downregulation, a drug that reverses the disease-associated gene expression changes may hold therapeutic potential for the disease. Genetically-anchored TWAS minimizes the problem of reverse causality. Indeed, Wu et al. applied PrediXcan models to find drug repurposing candidates using the Integrative Library of Integrated Network-based Cellular Signatures (iLINCS) platform(Wu et al., 2022). The latest release (CMAP LINCS 2020) of this platform contains over 3 million gene expression profiles with > 80,000 perturbations and > 200 cell lines. The resource is roughly a 3-fold expansion relative to the previous version (2017)(Subramanian et al., 2017). Methodologically, previous approaches (e.g., a widely adopted nonparametric pattern-matching method based on the Kolmogorov−Smirnov (KS) statistics(Lamb et al., 2006)) do not fully consider the pattern of alignment between the (transcript)omic features of drug induction and disease susceptibility, including, explicitly, genetically-determined risk, within a multitarget or polygenic framework. In summary, a genetics-informed omics-based repurposing framework that addresses critical methodological gaps is needed.

To address these gaps, here we developed Transcriptome-informed Reversal Distance (TReD) to integrate cell-based drug response profiles (CMAP LINCS 2020) and population-level disease signatures (TWAS and DGE) to identify drug repurposing candidates. For illustration, we applied the framework to search for drug repurposing candidates for COVID-19 and T2D.

## Results

### Study overview

Genetics-informed single-gene matching and omics profile-based signature matching are two promising approaches used in drug repurposing, with diverse types of omics data providing a valuable opportunity. In this study, we integrated these two approaches, leveraging transcriptome data to identify potential drug repurposing candidates (Figure 1). A single gene-based approach that combines TWAS and colocalization analysis (coloc-SuSiE) detects genes associated with disease (Fig 1A, Supplemental Fig. S1). Subsequently, drugs targeting these genes are prioritized as candidate therapeutics. For the multi gene-based approach, we developed TReD, integrating TWAS- and DGE- derived disease signatures and cell-based drug response profiles (CMAP LINCS 2020) (Fig. 1B, Supplemental Fig. S1). In this study, we focused primarily on the drug response profile data arising from small-molecule perturbations in immune-related cells and blood glucose metabolism-related tissues for the specific TReD applications to COVID-19 and T2D, respectively. TReD is based on transcriptome signature matching, which relies on the signature reversion principle (Wagner et al., 2015). In contrast to previous transcriptome signature pattern-matching studies, we proposed a “reversal distance” which quantifies the drug’s potential in reversing the disease-associated gene expression, using a metric that is scale-invariant and directionality-focused. To conduct hypothesis testing and evaluate the statistical significance, we employed a permutation approach.

**Figure 1.**
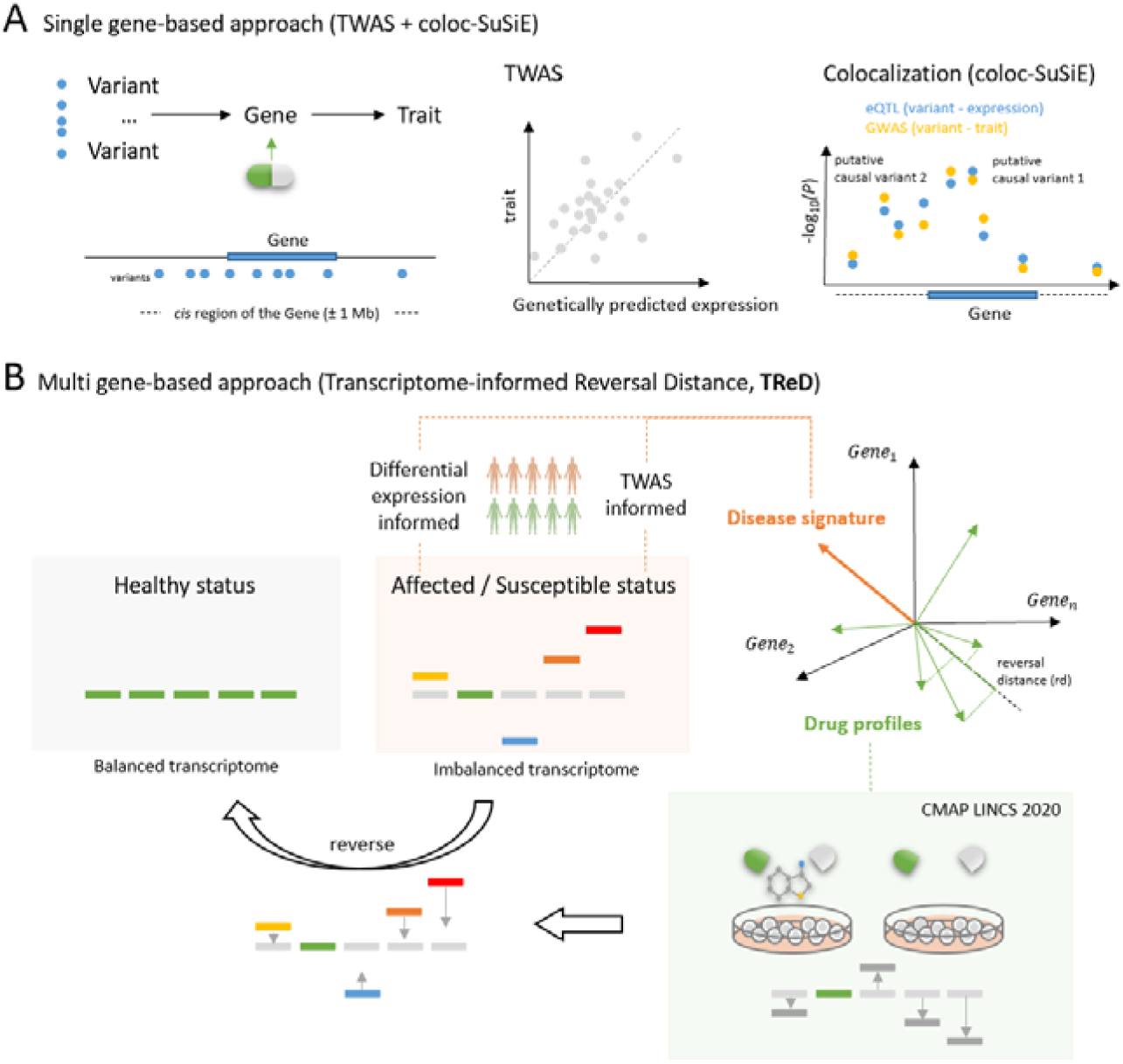
The framework overview. (A) Single gene-based approach integrates TWAS (e.g., PrediXcan) and colocalization analysis (e.g., coloc-SuSiE). (B) Multi-gene-based approach TReD (Transcriptome-informed Reversal Distance) utilizes DGE- and TWAS-derived population-level gene expression profiles as disease signatures, reflecting affection status and germline susceptibility, respectively. By embedding the population-level disease signatures (orange vector) and the cell-based drug response profiles (green vectors) in the same high-dimensional space, the reversal distance (*rd*) is estimated for each drug. A permutation test is used to estimate the statistical significance of the reversal distance (**Methods**).

### Single-target drug repositioning using TWAS and coloc-SuSiE

We performed TWAS to identify association between gene expression and a given phenotype. Models to predict gene expression were fit using a reference panel, subsequently applying gene expression models to GWAS of COVID-19 severity and T2D. We used data from Genotype-Tissue Expression (GTEx) v8, focusing on select tissues potentially relevant to COVID-19 severity (whole blood, lung, lymphocytes, and spleen samples(Vanderbeke et al., 2021)) or T2D (visceral adipose, liver, pancreas, skeletal muscle). Genes that survived Bonferroni correction for each tissue were considered significant (Fig. 2, Supplemental Table S1). For TWAS results significant after multiple testing, we performed coloc/coloc-SuSiE. eQTL and GWAS signals were determined to “colocalize” if maximum posterior probability of colocalization (i.e., PP.H4) was greater than or equal to 0.85 for coloc or coloc-SuSiE. The results of coloc are in Supplemental Table S2 and those of coloc-SuSiE are in Supplemental Table S3. We further examined which prioritized disease-related genes might be drug targets. Drug target genes were defined using public databases, including DrugBank(Wishart et al., 2018), the Drug Gene Interaction Database (DGIdb)(Freshour et al., 2021), and the Therapeutic Target Database (TTD)(Zhou et al., 2022).

**Figure 2.**
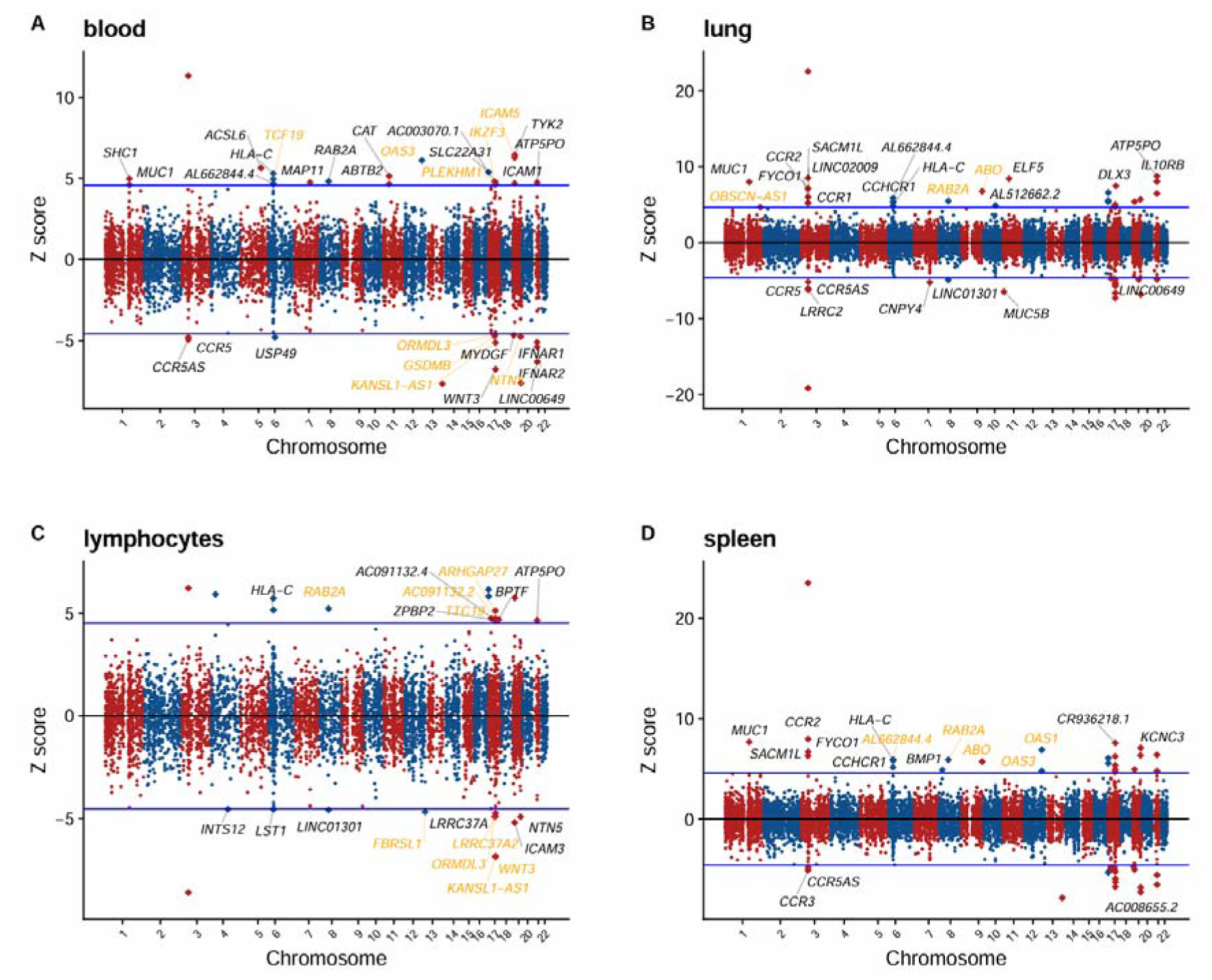
TWAS and colocalization analysis for COVID-19. Manhattan plots of TWAS results for COVID-19 severity in whole blood (A), lung (B), lymphocytes (C), and spleen (D), respectively. The *Z*-scores for TWAS are shown. *Z*-score > 0 indicates a gene for which higher genetically-determined expression is associated with COVID-19 severity. The labelled genes survived multiple testing corrections (over or below the blue lines) given the number of genes tested. Genes that further pass the significance threshold from the colocalization analysis are colored gold.

For COVID-19 severity, we identified 34, 45, 27, and 43 significant genes in blood, lung, lymphocytes, and spleen, respectively in TWAS. Nine genes showed significance across four tissues. Notably, *tyrosine kinase 2* (*TYK2)* was significant in all tissues with a concordant positive effect, signifying that an increased expression of *TYK2* would be associated with an increased risk of severe COVID-19. TYK2 is an intracellular kinase belonging to the Janus kinase (JAK) family, which plays a pivotal role in the Janus Kinase–Signal Transducer and Activator of Transcription (JAK-STAT) pathway(Hu et al., 2021). High expression of *TYK2* and host-driven inflammatory lung injury are related to life-threatening COVID-19(Pairo-Castineira et al., 2021). *RAB2A* also showed a positive effect in TWAS in the four tissues. *RAB2A* is required for SARS-CoV-2 replication(Li et al., 2020), and higher expression of *RAB2A* is associated with poorer COVID-19 outcome(Pairo-Castineira et al., 2023), consistent with directionality we observed.

In the four analyzed tissues, we identified a total of 28 unique genes post coloc/coloc-SuSiE (Supplemental Fig. S2). Among these targets, 50% (14/28) have been reported to be associated with COVID-19 (*DPP9*(Wang et al., 2021a), *WNT3*(Wu et al., 2021), *ABO*(Ellinghaus et al., 2020; Ray et al., 2021), *NAPSA*(Krishnamoorthy et al., 2023), *GSDMB*(Chang et al., 2022), *RAB2A*(Pairo-Castineira et al., 2023), *OAS1*(Banday et al., 2022), *OAS3*(Pairo-Castineira et al., 2021), *LRRC37A2*(Zhu et al., 2023), *MAPT*(Cao et al., 2023), *FUT2*(Andreakos et al., 2022; Kousathanas et al., 2022), *IKZF3*(Yao et al., 2023), *ICAM5*(Kousathanas et al., 2022), *CCR9*(Ellinghaus et al., 2020; Yao et al., 2021)). *CCR9* and *MAPT* are druggable genes: drugs such as docetaxel and astemizole, which target *MAPT,* had COVID-19-related support. Docetaxel can inhibit the active site of SARS-CoV-2 RdRP(Bagabir, 2023). Astemizole (antihistamine) can interfere with SARS-COV-2 entry (Wang et al., 2021b).

For T2D, 86 genes were identified by both TWAS and coloc/coloc-SuSiE in tissues examined (Fig. 3). We further investigated their associations with T2D in the Type 2 Diabetes Knowledge Portal(Costanzo et al., 2023). Among the 86 genes, ten have been classified as potential T2D effectors, including *Insulin receptor substrate 1* (*IRS1*), a gene central in insulin signal transmission(Sun et al., 1992). We also identified T2D-associated genes *PTH1R* and *NOTCH2*, both druggable. *PTH1R* showed a negative effect on T2D in visceral adipose tissue. Teriparatide (TPTD) and abaloparatide agonize *PTH1R,* and TPTD may improve glucose control in select populations (Mazziotti et al., 2014). Similarly, *NOTCH2* showed a negative effect on T2D in both liver and visceral adipose tissue (though drugs targeting NOTCH2 like nirogacestat and rg-4733 are inhibitory).

**Figure 3.**
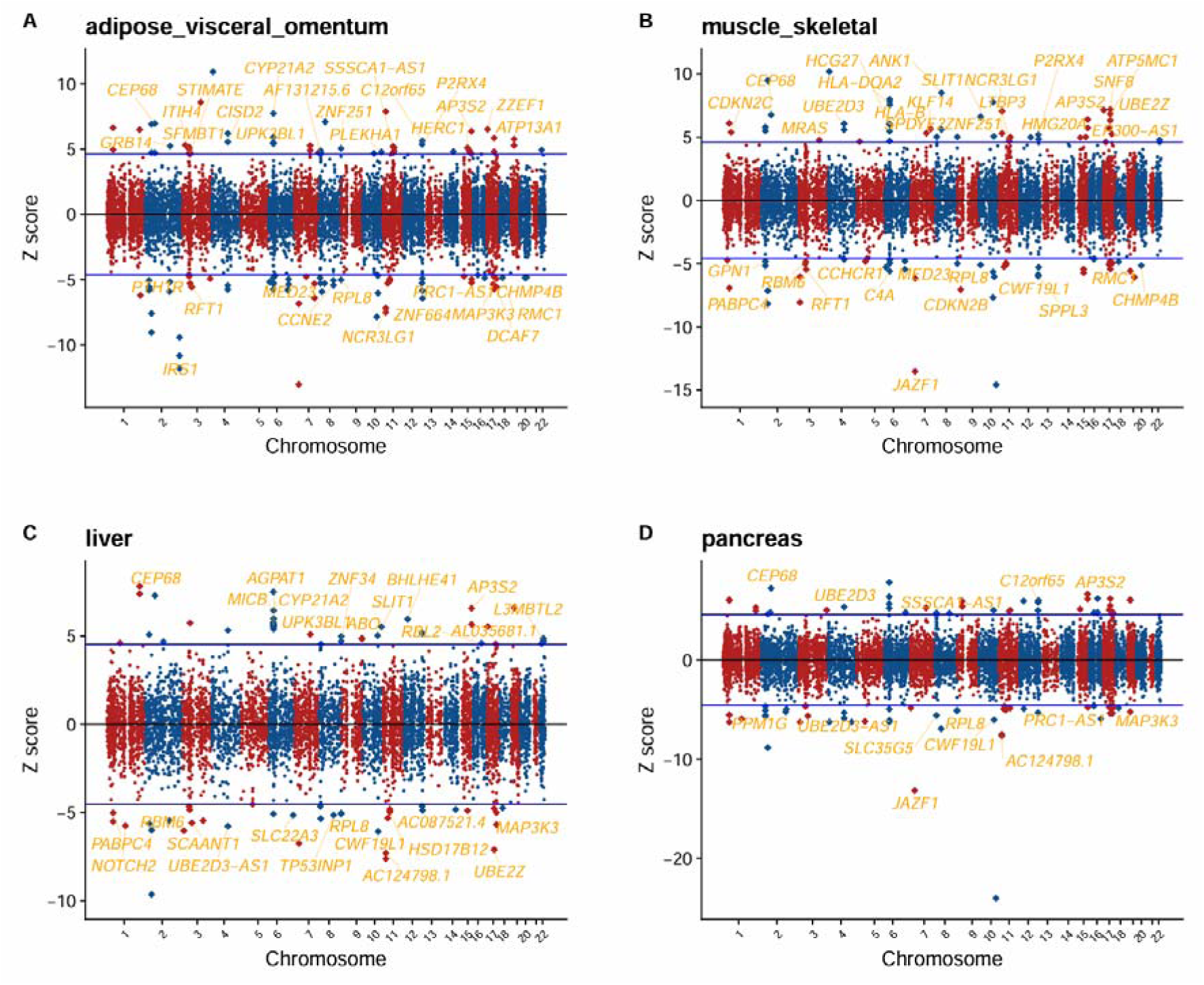
TWAS and colocalization analysis for T2D. Manhattan plots of TWAS results for T2D in visceral adipose (A), skeletal muscle (B), liver (C), and pancreas (D), respectively. The *Z*-scores for TWAS are shown. *Z*-score > 0 indicates a gene for which higher genetically-determined expression is associated with T2D. The labelled genes survived multiple testing corrections (over or below the blue lines) given the number of genes tested. Genes that further pass the significance threshold from the colocalization analysis are colored gold.

### Multi-target drug repositioning using TReD

Next, we applied TReD to prioritize potential drug repurposing candidates for COVID-19 and T2D. We sought to identify significant reversal effects from the cellular drug response profiles on the population-level disease signatures at transcriptome level (**Methods**). In addition to four TWAS-derived disease signatures, we included three DGE-based disease signatures (for COVID-19: ALV (alveolar), EXP (expansion), BALF (bronchoalveolar lavage fluid); for T2D: islets, myoblasts, myotubes; Table S4 and Methods). Rules for prioritizing drugs are in **Methods**. Briefly, we defined a metric called reversal distance (*rd*) as a metric of how a drug may counteract a disease signature (defined by cellular transcriptomic response to the drug). For a drug to be considered a potential candidate, its reversal distance must be two or more standard deviations away from the mean *rd* of the null distribution and show nominal significance in a permutation test. To ensure robustness, we further required consistent results across technical LINCS replicates, and at least one drug response profile analysis (permutation p-value) for the given drug must pass Bonferroni correction.

For COVID-19, we identified 707 drugs showing nominal significance in any of the seven disease signatures. For perturbations without repeated LINCS experiments, a consistent result in a confirmatory cell line is required to support the candidate drug response profile. Additionally, we highlighted drugs that exhibited a reverse effect in at least four disease signatures. Lastly, we prioritized 36 drugs for COVID-19 that show significance (*P_adj_* < 0.05) in at least one drug response profile (Figure 4). Table 1 provides an overview of the 36 drugs, including the reversal distance across the different disease signatures and whether there is any prior evidence linking to COVID-19 and/or clinical studies for the treatment of COVID-19. Notably, nearly 70% of the drugs (25/36) are reported related to COVID-19, and seven drugs have ongoing/completed clinical trials (Table 1).

**Figure 4.**
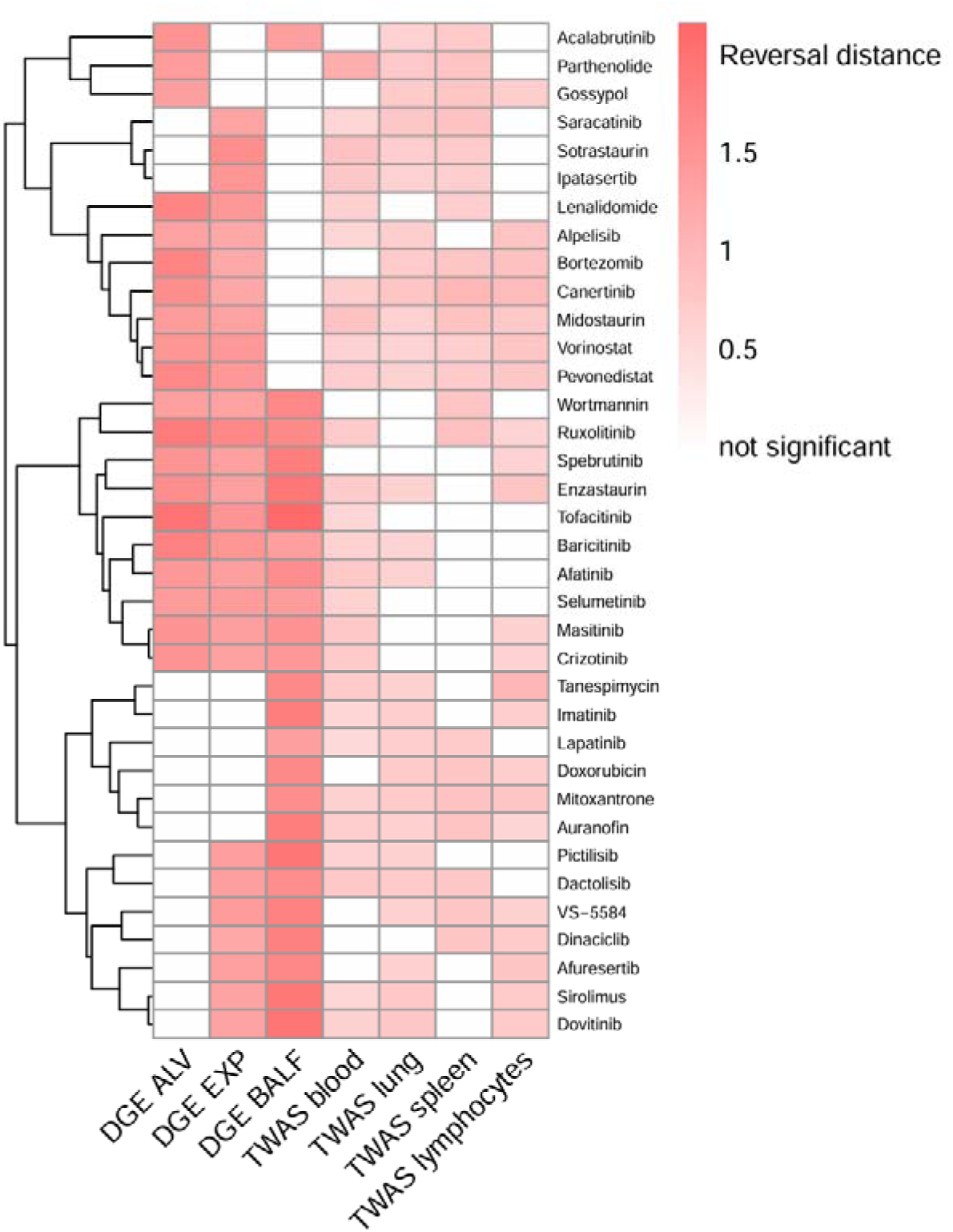
Drug repositioning candidates with potential reversal effect on COVID-19 disease signatures from DGE and TWAS. In total, 36 drugs were found by our Transcriptome-informed Reversal Distance (TReD) framework (**Methods**) to have a potential reversal effect on COVID-19 as indicated by at least four of the seven tested transcriptome-based disease signatures (x-axis). The heatmap shows the reversal distance for each drug response profile / disease signature pair. Blank cells denote “not significant” in the permutation test (the information on significance can be found in Supplemental Table S5). Hierarchical clustering was performed on the drug response profile (y-axis). The source data for the disease signatures, including TWAS (four tissue/cell types) and DGE (ALV, EXP, and BALF), are described in Methods.

**Table 1.** DrugBank annotations for drugs with significant reversal effects on at least four disease signatures for COVID-19.

| Drug Hit | Description | Reversal distance for each of the disease signatures |  |  |  |  |  |  | Evidence for COVID-19 | Evidence for clinical trials |
| --- | --- | --- | --- | --- | --- | --- | --- | --- | --- | --- |
|  |  | DGE | DGE | DGE | TWAS | TWAS | TWAS | TWAS |  |  |
|  |  | ALV | EXP | BALF | blood | lung | spleen | lymphocytes | 9 |  |
| Wortmannin | PI3Ks inhibitor | 1.350 | 1.311 | 1.699 | n.s. | n.s. | 0.829 | n.s. | Yes(Jiang et al., 2022) | No |
| VS-5584 | Antineoplastic | n.s. | 1.426 | 1.784 | n.s. | 0.644 | 0.794 | 0.640 | Yes(Bhara dwaj et al., 2022) | No |
| Vorinostat | Antineoplastic | 1.465 | 1.458 | n.s. | 0.632 | 0.597 | 0.700 | 0.813 | Yes(Sinha et al., 2020) | No |
| Tofacitinib | Antineoplastic and Immunomodulating | 1.991 | 1.518 | 2.159 | 0.583 | n.s. | n.s. | n.s. | Yes(Guim arães et al., 2021) | Yes |
| Tanespimycin | Antitumor antibiotics | n.s. | n.s. | 1.665 | 0.740 | 0.653 | n.s. | 1.026 | No | No |
| Spebrutinib | Immunomodulatory | 1.507 | 1.333 | 1.828 | n.s. | n.s. | n.s. | 0.634 | Yes(Kalia murthi et al., 2021) | No |
| Sotrastaurin | Immunosuppressant | n.s. | 1.625 | n.s. | 0.855 | 0.720 | 0.727 | n.s. | Yes(Huang et al., 2022) | No |
| Sirolimus | Immunosuppressant | 1.323 | 1.299 | 1.938 | 0.555 | 0.743 | n.s. | 0.733 | Yes(Bisch of et al., 2021) | Yes |
| Selumetinib | Antineoplastic | 1.426 | 1.406 | 1.411 | 0.667 | n.s. | n.s. | n.s. | Yes(Xie et al., 2022) | No |
| Ruxolitinib | Antineoplastic | 1.871 | 1.685 | 1.666 | 0.722 | n.s. | 0.878 | 0.592 | Yes(Lamontagne et al., 2020) | Yes |
| Pictilisib | PI3K inhibitors | n.s. | 1.377 | 1.944 | 0.615 | 0.636 | n.s. | n.s. | No | No |
| Pevonedistat | Antineoplastic | 1.686 | 1.450 | n.s. | 0.709 | 0.650 | 0.732 | 0.785 | No | No |
| Parthenolide | Anti-Inflammatory | 1.403 | n.s. | n.s. | 1.138 | 0.753 | 0.841 | n.s. | Yes(Bahra<br>mi et al.,<br>2020) | No |
| Mitoxantrone | Chemotherapy drugs | n.s. | 1.242 | 1.599 | 0.648 | 0.729 | 0.847 | 0.821 | Yes(Preet<br>et al.,<br>2022) | No |
| Midostaurin | Antineoplastic | 1.438 | 1.325 | n.s. | 0.863 | 0.672 | 0.853 | 0.748 | No | No |
| Masitinib | Antineoplastic and<br>Immunomodulating | 1.511 | 1.372 | 1.529 | 0.750 | n.s. | n.s. | 0.639 | Yes(Dray<br>man et al.,<br>2021) | Yes |
| Lenalidomide | Antineoplastic and<br>Immunomodulating | 1.750 | 1.441 | n.s. | 0.646 | n.s. | 0.714 | n.s. | Yes(von<br>Metzler et<br>al., 2022) | No |
| Lapatinib | Antineoplastic | n.s. | n.s. | 1.357 | 0.541 | 0.709 | 0.733 | n.s. | Yes(Saul<br>et al.,<br>2023) | No |
| Ipatasertib | Antineoplastic | n.s. | 1.478 | n.s. | 0.768 | 0.654 | 0.708 | n.s. | No | No |
| Imatinib | Antineoplastic | n.s. | n.s. | 1.814 | 0.552 | 0.700 | n.s. | 0.719 | Yes(van<br>de<br>Veerdonk<br>et al.,<br>2022) | Yes |
| Gossypol | Polyphenolic<br>compounds | 1.374 | n.s. | n.s. | n.s. | 0.738 | 0.824 | 0.711 | Yes(Wang<br>et al.,<br>2022) | No |
| Enzastaurin | Antineoplastic | 1.616 | 1.346 | 1.957 | 0.738 | 0.642 | n.s. | 0.840 | Yes(Huang et al., 2022) | No |
| Doxorubicin | Chemotherapy drugs | n.s. | n.s. | 1.660 | n.s. | 0.730 | 0.798 | 0.685 | Yes(Sajid Jamal et al., 2022) | No |
| Dovitinib | Multi-targeted kinase inhibitor | n.s. | 1.304 | 1.964 | 0.652 | 0.782 | n.s. | 0.737 | No | No |
| Dinaciclib | Antineoplastic | n.s. | 1.236 | 1.800 | n.s. | n.s. | 0.816 | 0.738 | Yes(Ahmad et al., 2023) | No |
| Dactolisib | Antineoplastic | n.s. | 1.363 | 1.629 | 0.761 | 0.732 | 0.772 | n.s. | Yes(Tomazou et al., 2021) | No |
| Crizotinib | Antineoplastic | 1.513 | 1.318 | 1.459 | 0.720 | n.s. | n.s. | 0.625 | Yes(Chen et al., 2021) | No |
| Canertinib | Antineoplastic | 1.613 | 1.253 | n.s. | 0.687 | 0.801 | 1.029 | 0.957 | No | No |
| Bortezomib | Antineoplastic | 1.766 | 1.241 | n.s. | n.s. | 0.734 | 0.819 | 0.883 | Yes(Li et al., 2022) | No |
| Baricitinib | Immunosuppressant | 1.767 | 1.493 | 1.393 | 0.633 | 0.594 | n.s. | n.s. | Yes(2022) | Yes |
| Auranofin | Antiinflammatory and Antirheumatic | n.s. | n.s. | 1.815 | 0.715 | 0.664 | 0.844 | 0.588 | Yes(Bijl et al., 2021) | No |
| Alpelisib | Antineoplastic | 1.319 | 1.249 | n.s. | 0.582 | 0.711 | n.s. | 0.840 | No | No |
| Afuresertib | Akt inhibitor | n.s. | 1.334 | 1.680 | n.s. | 0.669 | n.s. | 0.794 | No | No |
| Afatinib | Antineoplastic | 1.442 | 1.369 | 1.628 | 0.744 | 0.641 | n.s. | n.s. | No | No |
| Acalabrutinib | Antineoplastic | 1.526 | n.s. | 1.355 | n.s. | 0.650 | 0.722 | n.s. | Yes(2020) | Yes |
| Saracatinib | Antineoplastic | n.s. | 1.279 | n.s. | 0.576 | 0.771 | 0.859 | n.s. | No | NO |

The estimated reversal distance is shown. N.s. denotes not significant in the permutation test. The source data for the disease signatures, including TWAS (four tissue/cell types) and DGE (ALV, EXP, and BALF), are described in Methods.

**Table 2.**
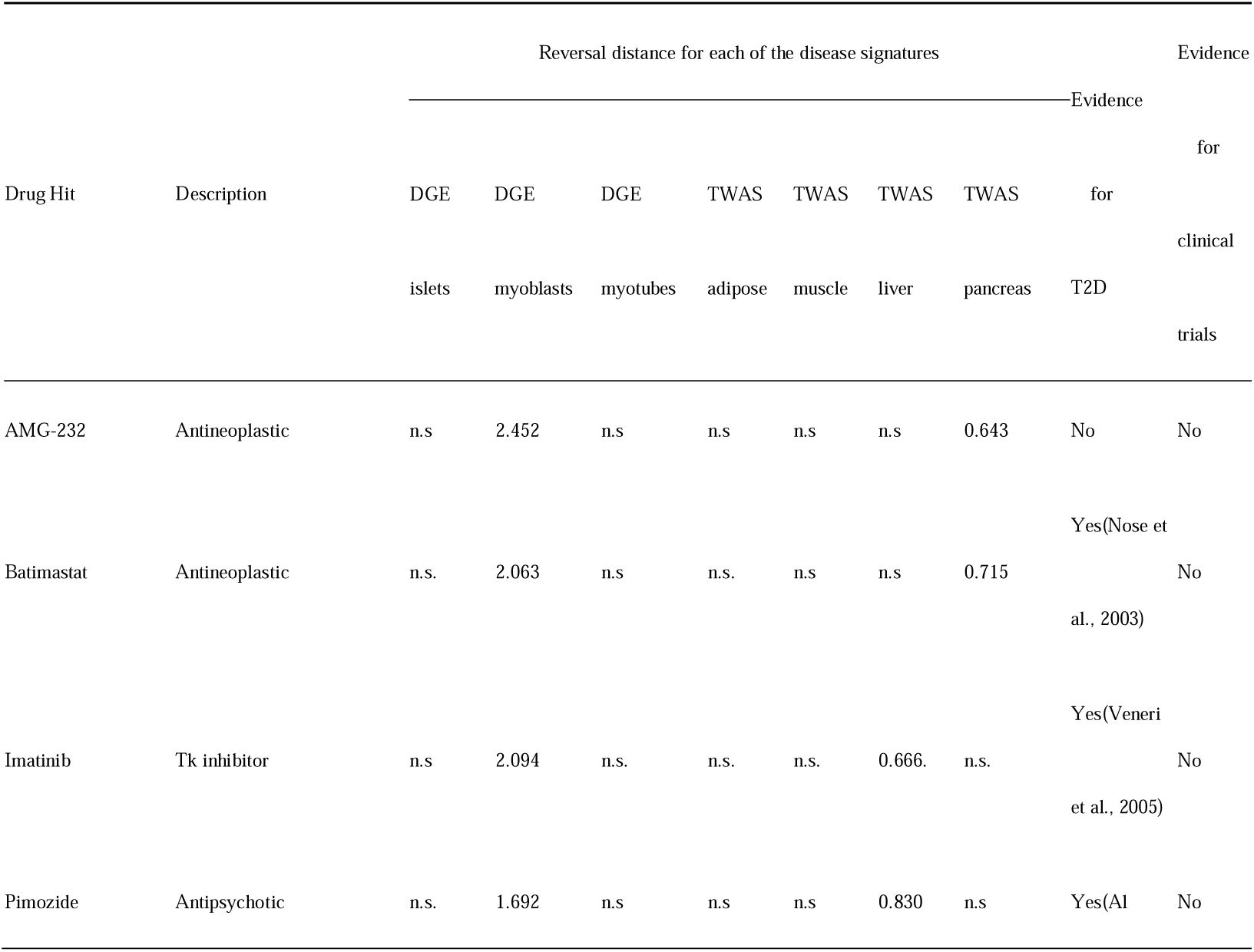
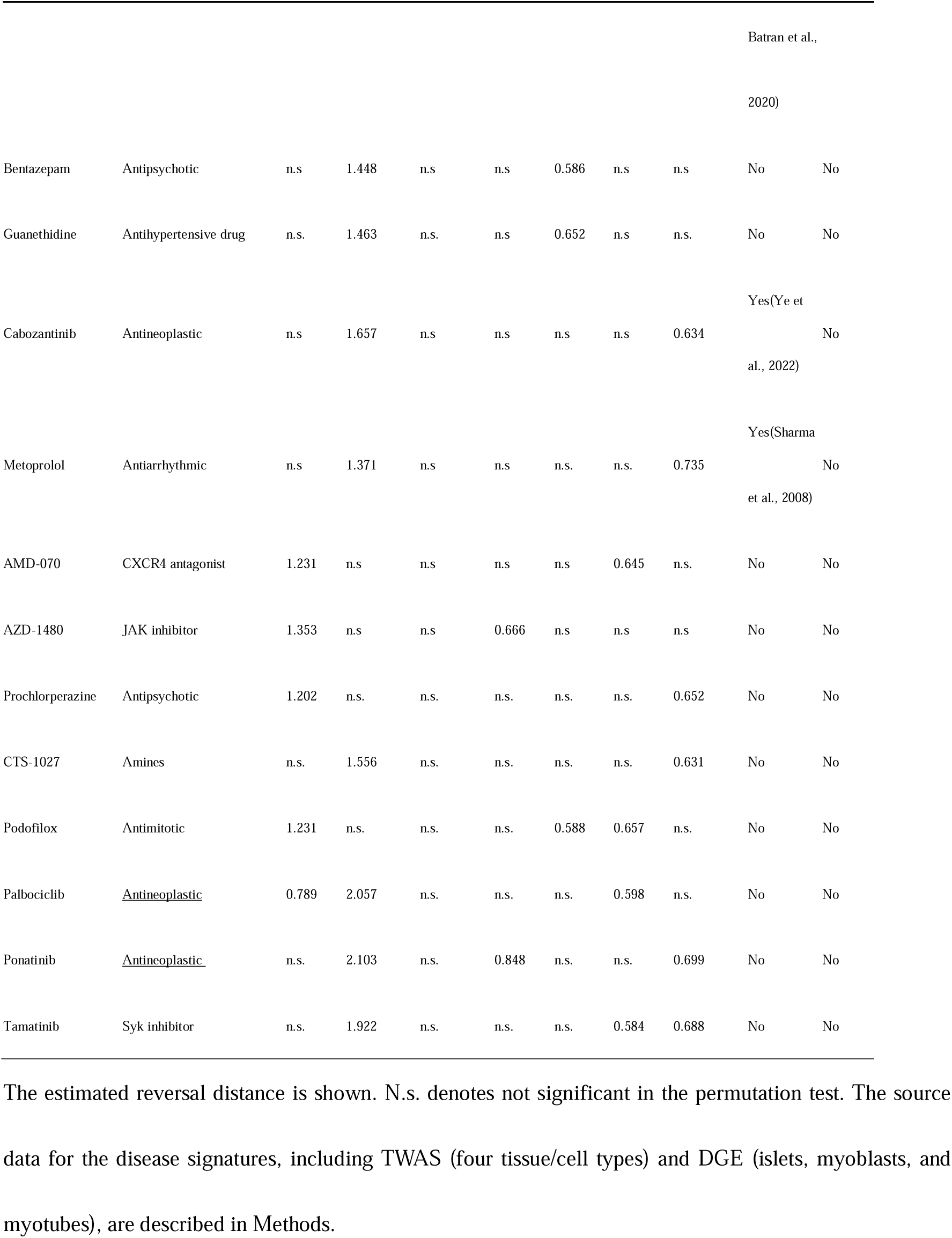
DrugBank annotations for drugs with significant reversal effects on two disease signatures for T2D.

Of note, over 40% of candidates are immunomodulatory drugs, especially JAK inhibitors. Baricitinib, significantly reversing five disease signatures (Fig. 4), is a JAK1/2 and TYK2 inhibitor approved for COVID-19 therapy in 2022. Ruxolitinib significantly reversed six disease signatures (Fig.4), with a variety of reports supporting divergent clinical efficacy (Koschmieder et al., 2020; Vannucchi et al., 2021) (Han et al., 2022) (Rein et al., 2022). Analogously, enzastaurin—investigational for treatment of glioblastoma multiforme via selectively inhibition of PKC-beta and other kinases involved in cell proliferation and survival—appeared effective in reversing six disease signatures (**Figure 4)**. Of note, SARS-CoV can activate the PKC members(Ji et al., 2009), and inhibitors of protein kinase C family members, including enzastaurin, may decrease SARS-CoV-2 replication *in vitro* (Huang et al., 2022)

For T2D, we identified 78 drugs/compounds as potential candidates for at least two T2D disease signatures (one TWAS and one DGE). We further filtered out 62 drugs/compounds without any drug response profile passing multiple comparison. This resulted in a total of 16 drugs/compounds (Fig. 5). Five of the drugs/compounds have been previously reported as potential options for the prevention or treatment of T2D. For example, imatinib (notably elevated reversal distance) has been shown to improve insulin sensitivity and maintain glucose homeostasis(Veneri et al., 2005) potentially via effects on endoplasmic reticula stress (Han et al., 2009). Pimozide also showed a significant reversal effect and may alleviate hyperglycemia in diet-induced obesity through inhibition of skeletal muscle ketone oxidation(Al Batran et al., 2020). Batimastat demonstrated a significant reversal effect, with potential roles in T2D-related complications as a matrix metalloproteinase inhibitor. (Ramnath et al., 2014). Collectively, these results suggest pathophysiologic plausibility of TReD-defined targets in T2D.

**Figure 5.**
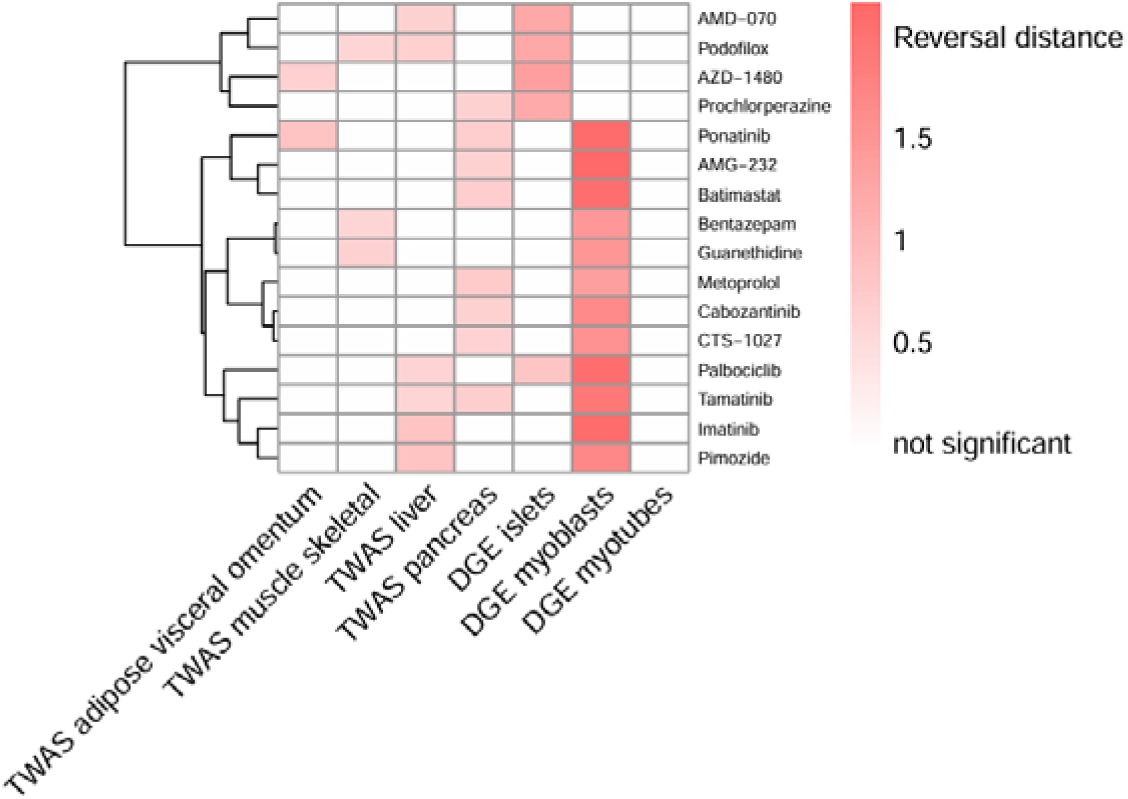
Drug repositioning candidates with potential reversal effect on T2D disease signatures from DGE and TWAS. In total, 16 drugs were found by our Transcriptome-informed Reversal Distance (TReD) framework (**Methods**) to have a potential reversal effect on T2D as indicated by two of the seven tested transcriptome-based disease signatures (x-axis). The heatmap shows the reversal distance for each drug response profile / disease signature pair. Blank cells denote “not significant” in the permutation test (the information on significance can be found in Supplemental Table S6). Hierarchical clustering was performed on the drug response profile (y-axis). The source data for the disease signatures, including TWAS (four tissue/cell types) and DGE, are described in Methods.

**Figure 6.**
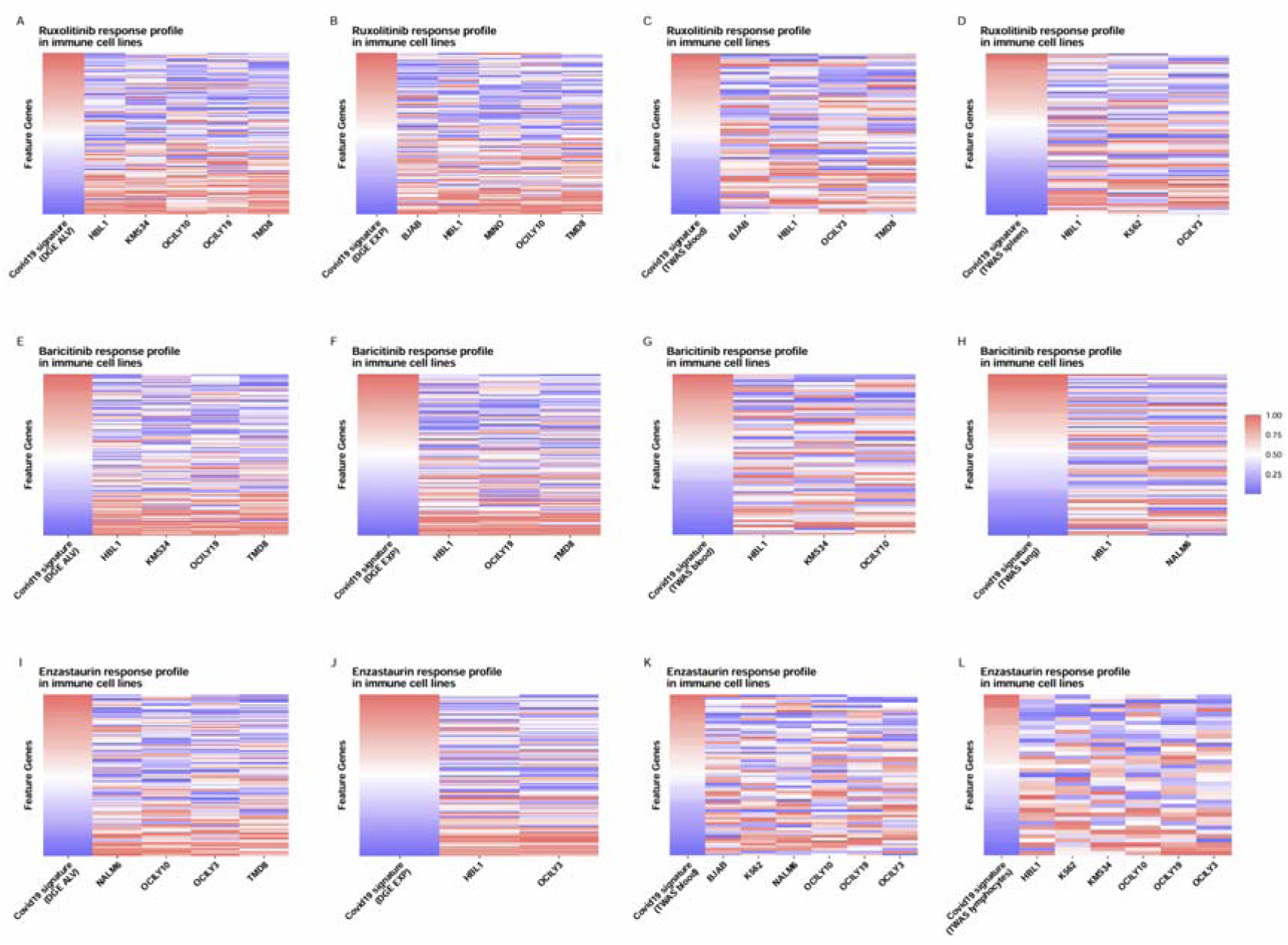
TReD-identified relationships between COVID-19 signatures and drug response profiles. The potential reversal effects of Ruxolitinib (A-D), Baricitinib (E-H), and Enzastaurin (I-L), which were identified against at least four disease signatures of COVID-19, are visualized using a heatmap. Taking (a) as an example, the first column in the heatmap represents differential gene expression (DGE) from the ALV gene expression disease signature ranked by fold change. Red and blue denote higher and lower expressed in affected samples, respectively. The remaining columns represent gene expression changes after Ruxolitinib treatment in immune cells. For drugs with multiple significant profiles, the drug response profiles (e.g., different duration after treatment) with the largest reversal distance are presented. The source data for the disease signatures, including TWAS (four tissue/cell types) and DGE (ALV and EXP), are described in Methods.

**Figure 7.**
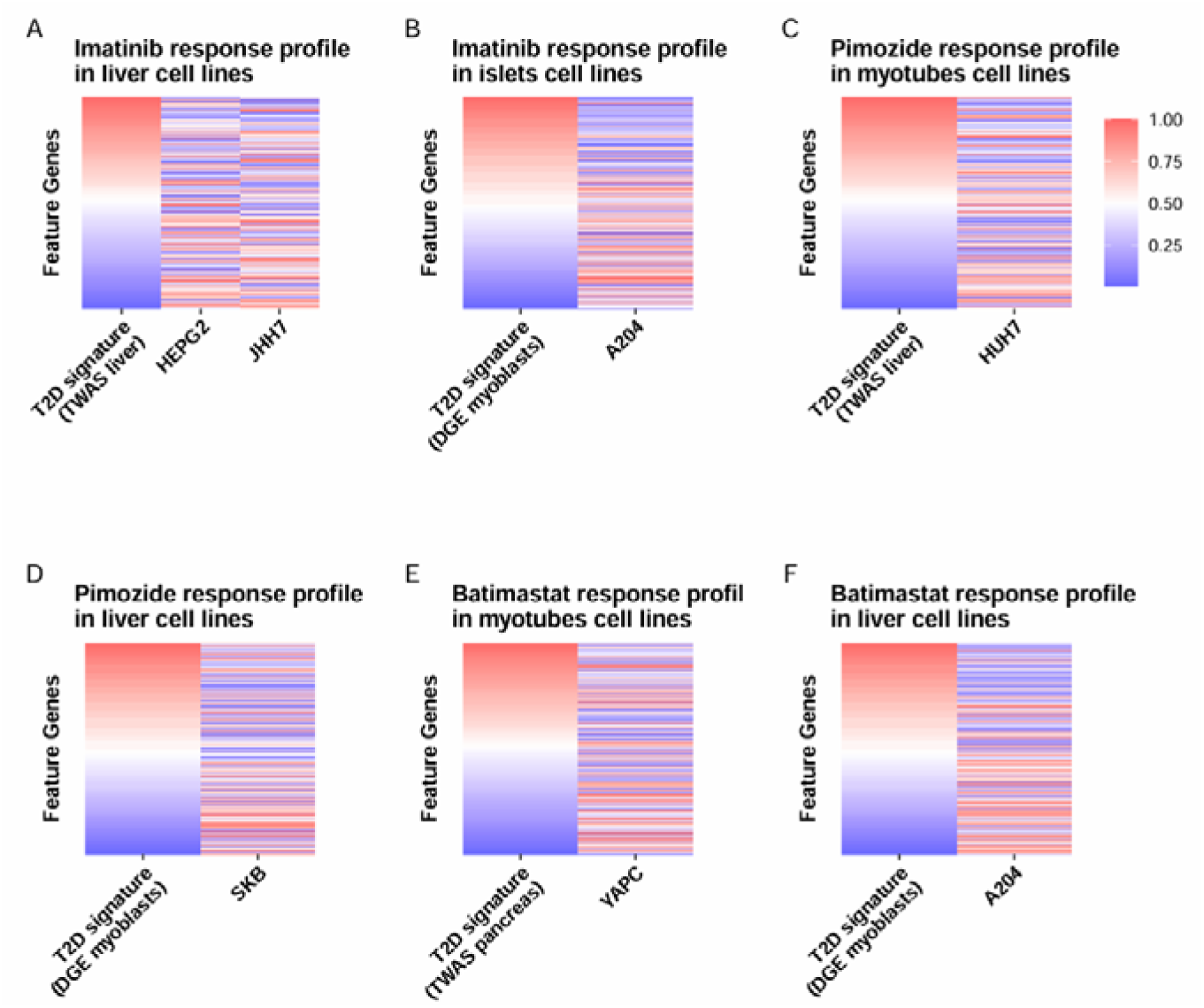
TReD-identified relationships between T2D signatures and drug response profiles. The potential reversal effects of Imatinib (A-B), Pimozide (C-D), and Batimastat (E-F), which were identified against two disease signatures of T2D, are visualized using a heatmap. Taking (a) as an example, the first column in the heatmap represents the transcriptome-wide association analysis (TWAS) derived disease signature in liver ranked by fold change. Red and blue denote higher and lower expressed in affected samples, respectively. The remaining columns represent gene expression changes after drug treatment in liver cells. For drugs with multiple significant profiles (e.g., different duration after treatment), the drug response profiles with the largest reversal distance are presented. The source data for the disease signatures, including TWAS (four tissue/cell types) and DGE, are described in **Methods**.

## Discussion

Drug development pipelines have traditionally relied on high-throughput rational or biologic synthesis and screening approaches. Recent high-dimensional genomic and transcriptional approaches have yielded a path to prioritize potential targets for directed study(Sass et al., 2024). However powerful, traditional genomic approaches (e.g., TWAS) alone only identify potential pathways to target, without taking advantage of the large bank of information on druggability and cellular responses to drugs already in existence. Here, we developed a framework to identify potential drug repurposing candidates by combining population-level disease signatures via human genetics (TWAS/DGE) and cellular responses to drugs at the transcriptional level (TReD). These approaches were successfully applied to COVID-19 and T2D, identifying known (e.g., baricitinib/TYK2) and novel drug-disease combinations.

Using TReD, we found a significant correlation between potential drug candidates and associated disease, specifically within pathophysiologies central to the two conditions studied. For example, in COVID-19, nearly half (16/36) of identified drugs were immunomodulatory (critical to COVID-19 pathogenesis), with prior study in COVID-19. Indeed, one target identified by TReD has in-human approval as a COVID-19 therapeutic (baricitinib in severe COVID-19), with others recommended as an alternative (tofacitinib (2021)). Our results in T2D were similar in scope. Among the prioritized drugs, we have highlighted pimozide and imatinib as having strong associations with T2D. Collectively, these results demonstrate the ability of this novel TReD framework to identify drug repositioning candidates.

We also investigated the drugs that are *positively* correlated with the disease signatures of COVID-19. A few such top-ranked drugs (including testosterone(Yassin et al., 2023) and otamixaban(Hempel et al., 2021)) were also supported by published studies as potential treatment options. This scenario may arise, for example, when a disease signature targeted by such a drug lies in a co-expression network with another disease signature whose net reversal is a potential therapeutic option. This result indicates that, in reality, the biological system operates in a much more complex fashion.

The strengths of our framework stem from several key aspects. Firstly, the amalgamation of single-target and multiple-target approaches enables a more comprehensive therapeutic search, and this is particularly valuable for addressing complex diseases involving multiple biomolecules or mechanisms. Secondly, the framework incorporates two complementary methods (DGE and TWAS) as population-level sources for disease signatures. Although TWAS may be underpowered with a limited sample size, a critical feature of such a genetics-anchored method ensures that it identifies *potentially* regulatory or candidate causal genes for the disease, rather than genes altered by the disease. In contrast, DGE is prone to identify the transcriptomic consequences of the disease; nevertheless, it remains well-powered even with limited sample sizes. The combination of these two methods should enrich for upstream mediators and molecular consequences, facilitating the discovery of new drug indications or the repositioning of existing drugs. Thirdly, the use of the reversal distance measurement in our study capitalizes on available (and growing) information from transcriptome-level disease-associated gene features and drug response profiles, with biologically relevant interpretation of an effective therapeutic effect on disease. This differs from previous and, we would argue, less interpretable methods, such as those that have relied solely on a non-parametric KS statistic, focusing on the maximum difference in the relative ranks of the upregulated and downregulated genes. Fourthly, we leveraged the latest CMAP LINCS 2020 data which contains over three million gene expression profiles with > 80,000 perturbations and > 200 cell lines affecting over 12,328 genes, providing a comprehensive candidate pool for drug repositioning. While our study focused on COVID-19 and T2D, our framework is highly generalizable and can be implemented to identify drug repositioning candidates for other diseases.

While encouraging as a first attempt to integrate TWAS and cellular response profiles, TReD has a key limitation: this approach is dependent on comprehensive curation of drug response profiles over a wide range of drug targets, cell types, and disease conditions. Indeed, while we were broad in scope for inclusion of curated data for these elements, the cellular response profiles were obtained primarily in cancer cell lines, which may not necessarily phenocopy the cell types and biological context of interest for the condition under study (e.g., COVID-19). Although we requested a confirmatory cell line with concordant response to support the findings, the treatment dose and duration in the confirmatory cell line are not always consistent. In addition, we recognize that most medications here are necessarily anti-cancer therapeutics, which may not be clinically translatable to other broad conditions (e.g., T2D) due to off-target (side) effects. Finally, other “omes” beyond transcriptional responses are likely relevant (e.g., protein expression, non-coding RNA) that may encode trans-organ signaling pathways and a broad cell atlas relevant to multi-system disorders (e.g., T2D). Our study does not consider the complex issues of drug-target interactions and drug-drug interaction, and consideration of in vivo pharmacokinetics, pharmacodynamics, and substrate processing are critical.

The solution to these limitations is either a strength of older drug discovery approaches—broad, rapid screening functional assays across cellular phenotypes and broad synthetic or derivatized chemical libraries or biologics for screening (e.g., rapid lentiviral CRISPR screening(Chaffin et al., 2022))—or a strength of ongoing drug development pipelines (Phase 0 and animal system studies). The role for TReD in this pipeline is to hone the potential space of therapeutics and provide a multi-dimensional platform to integrate across molecular genetic information to improve drug discovery efficiency.

In summary, integrating transcriptome-level profiles from cell-based drug response profiles and population-based disease signature studies, we developed a framework, TReD, to identify drug repurposing candidates. Deploying TReD in COVID-19 and T2D successfully identified both known and novel candidates that capture underlying pathobiological mechanisms important in each disease. These results suggest that computational approaches that unite molecular genetic information with cellular response profiles may be effective in drug repositioning across a wide array of disorders. Future work collating cell types, pharmacotherapies, and disease states is needed to fully realize the potential of this and analogous approaches.

## Methods

### GWAS for COVID-19 severity

The GWAS summary statistics for COVID-19 severity were downloaded from COVID-19 Host Genetics Initiative (https://www.covid19hg.org/), the latest version, release 7 of the hospitalized versus population data. The hospitalized phenotype refers to hospitalization with a laboratory confirmation of SARS-CoV2 infection. This dataset contains 44,989 cases and 2,356,386 controls. The summary statistics of GWAS of T2D were obtained from Xue et al’s study. This dataset contains 62,892 T2D cases and 596,424 controls of European ancestry(Xue et al., 2018). (http://cnsgenomics.com/data/t2d/Xue_et_al_T2D_META_Nat_Commun_2018.gz). Additional descriptions of the included studies can be found in Supplemental Table S7.

### TWAS and colocalization for COVID-19 severity and T2D

To identify disease associated genes, we applied TWAS with the best prediction models among PrediXcan(Gamazon et al., 2015), UTMOST(Hu et al., 2019), and JTI(Zhou et al., 2020) according to the prediction accuracy estimated by 5-fold cross-validation. We used the residuals of the normalized gene expression after regressing out relevant covariates. SNPs in the cis region (i.e.,1 Mb upstream and downstream) of the gene were considered candidate features for the gene’s prediction models. Applying the prediction model to GWAS dataset, we would estimate the genetically predicted expression level for each individual. The significance of the effect of the predicted expression of a gene on a trait was determined by linear regression. In practice, summary statistics-based association test was performed. Summary-statistics based association test utilizes the weights from the prediction model of the gene under consideration, the covariances of the SNPs included in the prediction model for the gene, and the GWAS effect size coefficient for each SNP. Notably, the summary statistics-based approach results in nearly perfect concordance with the individual-level based approach(Barbeira et al., 2018).

SARS-CoV-2 directly attacks pulmonary alveolar cells and causes lung injury. Pulmonary infections leading to COVID-19 induce adaptive immune responses while effector immune cells are generated and recruited in the lymphocytes, spleen, and blood. Therefore, blood, lung, lymphocytes, and spleen were considered as relevant tissues for COVID-19. Visceral adipose, liver, pancreas, and skeletal muscle were considered relevant tissues for T2D. For TWAS results that passed Bonferroni multiple testing correction, we performed colocalization to provide additional support under different sets of assumptions. To perform colocalization analysis in the presence of multiple potentially causal variants, we applied coloc(Giambartolomei et al., 2014) to the GWAS hits decomposed by SuSiE(Wallace, 2021) in each GWAS locus, which spans the SNPs located within 200 kb (in each direction) of the lead SNP. Notably, *coloc-SuSiE* is more generalizable since it relaxes the “one causal variant” assumption of regular *coloc*. We used European individuals in the 1000 Genomes Project as the LD panel. If SuSiE failed to converge after 1000 iterations for either eQTL or GWAS summary statistics, we instead used the regular *coloc*. eQTL and GWAS signals were determined to “colocalize” if the maximum posterior probability of colocalization (i.e., PP.H4) was greater than or equal to 0.85 using either *coloc* or *coloc-SuSiE*. TWAS significant genes showing positive colocalization evidence were then used to search for drugs targeting the genes, via DrugBank, TTD, DGIdb. DrugBank is a comprehensive and up-to-date resource for searching over 500,000 drugs and drug products, their targets, pathways, indications, and pharmacology. TTD is a comprehensive database of therapeutic targets, drugs, biomarkers and scaffolds for various diseases and ICD identifiers. DGIdb is an online resource on drug-gene interactions and the druggable genome(Cannon et al., 2024).

### DGE for COVID19 and T2D

In COVID-19, the DGE data—consisting of independent gene expression signatures labelled ‘ALV’, ‘EXP’, ‘BALF’—were derived from the study of Le et al(Le et al., 2021). The ALV signature was generated from RNA-seq data in human adenocarcinomic alveolar basal epithelial cells (GEO Dataset: GSE147507; n=67) (Blanco-Melo et al., 2020) (Supplemental Table S8). After Benjamini-Hochberg adjustment, the study identified 120 differentially expressed genes — 100 upregulated and 20 downregulated. The EXP signature was obtained by RNA-seq on organoid samples (Lamers et al., 2020) (Supplemental Table S9) (GSE149312; n=22). After Benjamini-Hochberg adjustment and applying a fold-change cutoff (|log2 FC| >I_2), 125 genes remained significant. The BALF signature was estimated by RNA-seq analysis of the bronchoalveolar lavage fluid samples from two COVID-19 patients and three controls (Xiong et al., 2020). After DESeq2 processing, 1349 genes were used to define the BALF signature (Table S10).

Three DGE datasets (labelled as ‘islets’, ‘myoblasts’, and ‘myotubes’) were included in the T2D analysis. The islet signature was generated from microarray data on pancreatic islets derived from T2D cases and normal glucose-tolerant controls (Gunton et al., 2005), consisting of 370 differentially expressed genes at p < 0.01 (Supplemental Table S11). The other two signatures were obtained from the Davegårdh C et al study (Davegårdh et al., 2021). Based on a false discovery rate less than 5%, the study identified 577 differentially expressed genes in myoblasts and 42 differentially expressed genes in myotubes (Supplemental Table S12 and Table S13)

### The CMAP LINCS 2020 dataset

The Library of Integrated Network-Based Cellular Signatures (LINCS) is a NIH Common Fund program. Its primary goal is to generate a comprehensive atlas of perturbation-response signatures. The CMAP LINCS 2020 data contains drug response profiles (defined by drug-induced gene expression profiles in response to perturbations). Here, we mainly focused on the transcriptomic response to small-molecule (drugs/compounds) perturbations in immune-related cells and blood glucose metabolism-related tissues for the application of the TReD to COVID-19 and T2D, respectively. We downloaded the level 5 beta data consisting of drug response profiles arising from small-molecule perturbations from CMAP LINCS 2020 (https://s3.amazonaws.com/macchiato.clue.io/builds/LINCS2020/level5/level5_beta_trt_cp_n720216x12328.gctx),

### Candidate drug prioritization

To quantify the extent of reversal on disease of a specific drug, we used the cosine similarity metric. For a given disease, we first represent the set of disease signature as a vector in a high-dimensional space (whose dimension *n* is the total number of genes) with each nonzero element of the vector given by a disease-associated gene. The disease signature can be derived from either TWAS or DGE. The vector for each candidate drug is embedded in the same *n*-dimensional space.

For each disease signature vector and drug response profile pair, we compute the reversal distance. Reversal distance measures a drug’s ability to counteract disease-related gene expression. A positive value suggests potential preventive or therapeutic effects, while a negative value indicates the drug may worsen the disease. A higher positive reversal distance suggests a stronger counteracting effect.

Let 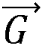 denote a disease signature vector, 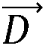 a drug profile vector, and *rd* the reversal distance. The *rd* quantifies the potential effect of the drug in reversing the disease signatures:

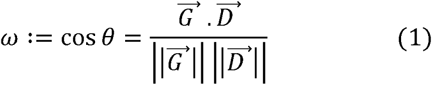

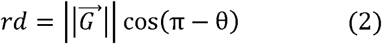

Here, 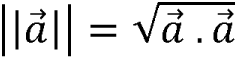 denotes the Euclidean norm of the vector 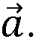 Note that the cosine similarity *ω* (equation 1) always lies in [-1,1], with-1 indicating exactly opposite orientation of the disease signature and the drug response profile and +1 indicating parallel orientation. The change in *ω* in response to a perturbation in the *i*-th element of the drug response profile 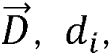 or the *i*-th element of the gene signature 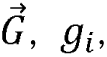 is given by:

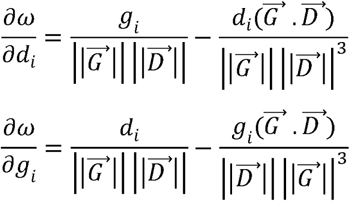

Each quantity measures the sensitivity of the metric. In equation 2, θ, and thus the reversal distance *rd*, is a function of 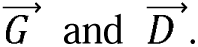 When we wish to emphasize this dependence, we will write *rd* as 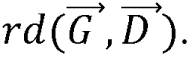. Equation 2 defines a function:

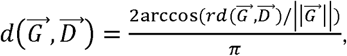

which satisfies the conditions for a distance function, namely, non-negativity: 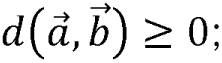 symmetry: 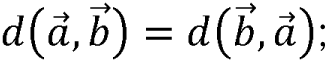 the triangle inequality; and the coincidence axiom: 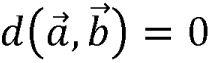if and only if 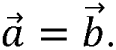 Despite the symmetry of the distance function, there is clearly an underlying *asymmetry*, as the disease signature vector 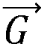 determines which genes are the “relevant” elements of the drug response profile vector 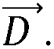

Of interest to us is the sampling distribution of the cosine similarity *ω*. Consider the function 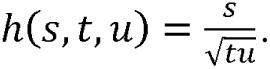 Note that *ω* is equal to 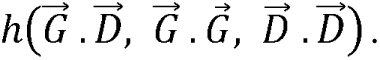 Furthermore, 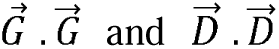 are 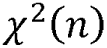 distributed. We leverage the so-called Delta Method. Using the first two terms of the Taylor series expansion and setting 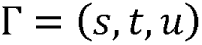 with expected value 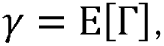 we obtain approximate expressions for the mean and variance of 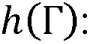

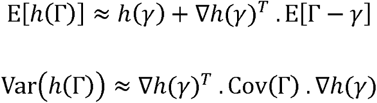

where 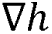 is the gradient.

In this framework, drug repurposing is a search problem: the vector 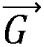 is evaluated against all possible 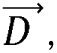 which are prioritized using *rd*. A larger *rd* indicates a stronger reversal effect. Assuming *K* total number of drugs tested and the reversal distance for the *i*-th drug, 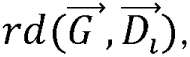 we get a probability distribution consisting of the *K* probabilities *p_i_* conditional on the disease signature information:

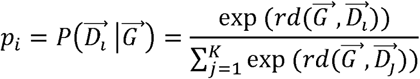

by applying the softmax function. One drug repurposing approach is to identify the drug that maximizes the reversal distance:

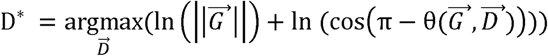

Here, we instead prioritized a potentially larger set of drugs, i.e., the top candidates that meet a significance threshold.

We converted the gene-disease association signals into rank ratios, which, for each gene, is defined as the rank of the gene divided by the total number of genes. Similarly, each element of the drug response profile was rank-transformed.

For each drug response profile, the p-value of the reverse distance (*rd*) is estimated by a permutation test. To generate the null distribution, we randomly shuffle the labels 1000 times to create numerous permuted samples. These samples are generated under the assumption of the null hypothesis, meaning there is no effect or association. In practice, the rank-ratio of disease signature is shuffled randomly. A null reversal distance is calculated on the basis of the shuffled disease signature. Then, we calculate the real reversal distance and estimate the proportion of permuted reversal distance that are as extreme or more extreme than the reversal distance obtained from the real data. This proportion represents the p-value. For each of the disease signatures, a significant drug response profile is defined as showing nominal significance in the permutation test (*P* < 0.05) and large *rd* (over 2 standard deviations from the mean across all the drug response profiles). In addition, we further adjust the experiment-wise permutation p-value (pooling different perturbations) across the tested cell lines using Bonferroni correction. For each disease, drugs with at least one significant (*P_adj_* < 0.05) drug response profile were prioritized.

We also implemented a degree-preserving permutation approach. Let *G* = (*V*, *E*) be a graph, where *v* is the set of nodes (genes) and *E* is the set of edges (defined by a co-expression network in a given tissue). To generate a permutation which preserves the degree distribution of the network, we randomly select two edges (say, *a* ↔ *b* and *c* ↔ *d*) with non-overlapping nodes, remove the edges between the nodes, and create new edges (*a* ↔ *c* and *b* ↔ *d*) by swapping the nodes until a certain proportion (80%) of the edges have been shuffled. The null reversal distance is computed for each such permutation.

### Application of TReD to COVID-19 severity and T2D

Leveraging the TWAS/DGE results and the cell-based gene expression data from CMAP LINCS 2020, we implemented the TReD framework to prioritize drug repositioning candidates for COVID-19 and T2D. Immune-related and T2D-related cell lines (matching the T2D-related tissues) have been annotated by multiple resources including the CMAP LINCS 2020 and the CCLE (cancer cell lines which have been used to study COVID-19 and other diseases(Bakowski et al., 2021)). Information on drugs and drug targets is curated in the bioinformatics and chemoinformatics resource DrugBank (<u>DrugBank Release Version 5.1.10 | DrugBank Online</u>) A total of 1112 drugs and 12 immune-related cell types were included for COVID-19, while 1980 drugs and 10 related cell types were included for T2D.

We leveraged the TWAS results of multiple tissues and three DGE profiles as disease signatures (as defined in the previous section). For perturbations with repeated LINCS experiments, consistent direction of *rd* is required. i.e., drug response profile with repeated experiments showing inconsistent direction of *rd* will be removed. Considering the multiple testing, we prioritize drugs which show at least one significant drug response profile (*P_adj_* < 0.05) for each disease.

### Software Availability

The source codes used in this study are available at GitHub(https://github.com/zdangm/TReD). Source codes are also provided as Supplemental Code.

## Acknowledgments

This research is supported by the National Natural Sciences Foundation of China 82204118 (D.Z.), the Key Laboratory of Intelligent Preventive Medicine of Zhejiang Province 2020E10004 (D.Z.), the National Institutes of Health (NIH) Genomic Innovator Award R35HG010718 (E.R.G.), NIH/NHGRI R01HG011138 (E.R.G.), NIH/NIA R56AG068026 (E.R.G.), and NIH/NIGMS R01GM140287 (E.R.G.). R.V.S. is supported by grants from the National Institutes of Health.

## Author contributions

Conceptualization, Dan Zhou and Eric R. Gamazon; formal analysis, Chunfeng He, Yue Xu, Yuan Zhou, and Ran Meng; writing--original draft, Chunfeng He, and Yue Xu; writing--review & editing, Dan Zhou, Eric R. Gamazon, Yuan Zhou, Jiayao Fan, Chunxiao Cheng, Lang Wu and Ravi V. Shah.

