## supplemental file 1 (Figure S1-S2) for "Integrating population-level and cell-based signatures for drug repositioning": supplemental file 1 (Figure S1-S2).docx

**Supplementary files**

**Supplemental Figure S1**: Application to identify drug repurposing candidates for COVID-19

**Supplemental Figure S2:** Venn diagram of TWAS significant and colocalization positive genes in four COVID-19 relevant tissues (blood, lung, lymphocytes, spleen)

**Supplemental Figure S1**: **Application to identify** **drug repurposing candidates for COVID-19** (a) GWAS summary statistics of COVID-19 severity and eQTL of four immune-related tissues were combined by PrediXcan and coloc/coloc-SuSiE, respectively. Genes that were both significant in TWAS after Bonferroni correction and exhibited positive colocalization signals (PP.H4 for coloc and coloc-SuSiE ≥ 0.85) were searched as potential drug targets. (b) Four TWAS results (*P_FDR_* < 0.05), three DGE for COVID-19 severity and cellular-level transcriptome drug profiles from CMAP LINCS 2020 were integrated by TReD to obtain drugs that reverse disease signature.

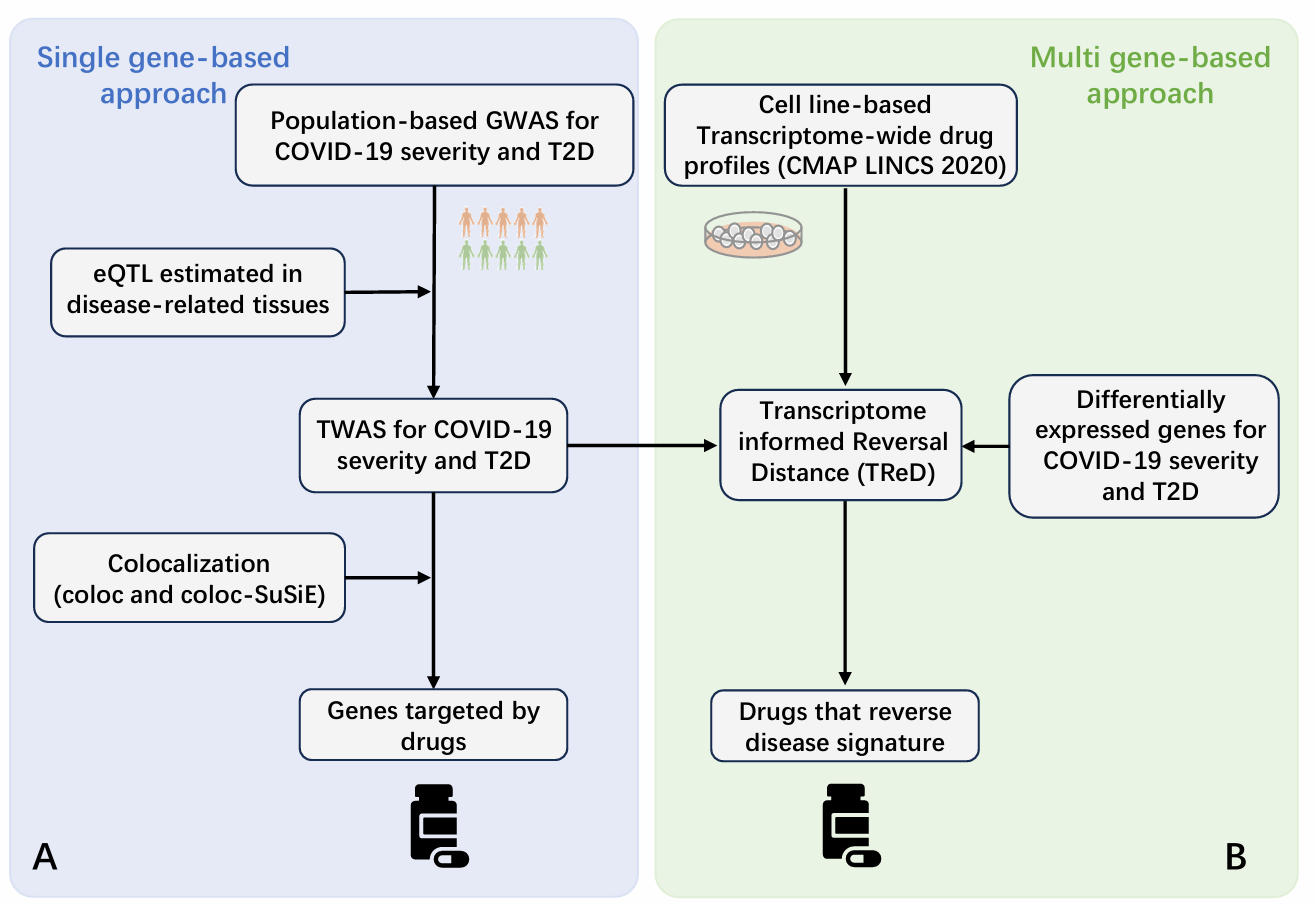

**Supplemental Figure S2: Venn diagram of TWAS significant and colocalization positive genes in four COVID-19 relevant tissues (blood, lung, lymphocytes, spleen)**. Blood had eleven genes, lungs had twelve, lymphocytes carried ten, and the spleen hosted twelve genes. In total, there were 28 distinct genes and eleven genes appeared in at least two tissues. Two genes overlapped among the four tissues.

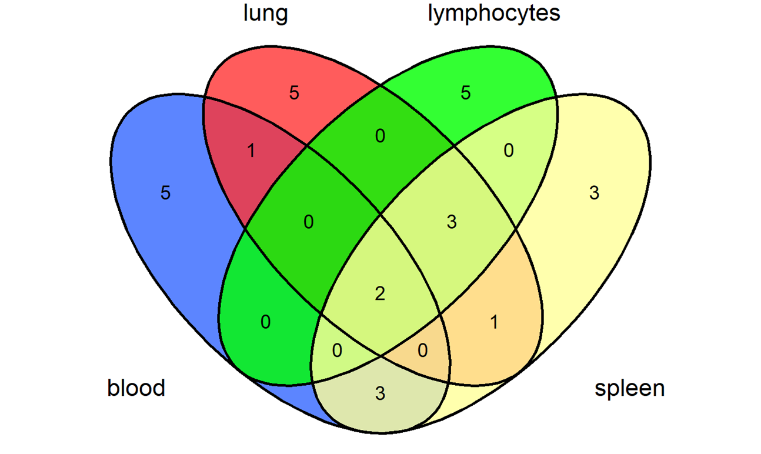
